# Evidence for “inter- and intraspecific horizontal genetic transfers” between anciently asexual bdelloid rotifers is explained by cross-contamination

**DOI:** 10.1101/150490

**Authors:** Christopher G. Wilson, Reuben W. Nowell, Timothy G. Barraclough

## Abstract

Bdelloid rotifers are microscopic invertebrates thought to have evolved for millions of years without sexual reproduction. They have attracted the attention of biologists puzzled by the maintenance of sex among nearly all other eukaryotes. Bdelloid genomes have an unusually high proportion of horizontally acquired non-metazoan genes. This well-substantiated finding has invited speculation that homologous horizontal transfer between rotifers also may occur, perhaps even 'replacing' sex. A 2016 study in *Current Biology* claimed to supply evidence for this hypothesis. The authors sampled rotifers of the genus *Adineta* from natural populations and sequenced one mitochondrial and four nuclear loci. For several samples, species assignments were incongruent among loci, which the authors interpreted as evidence of "interspecific genetic exchanges". Here, we use sequencing chromatograms supplied by the authors to demonstrate that samples treated as individuals actually contained two or more divergent mitochondrial and ribosomal sequences, indicating contamination with DNA from additional animals belonging to the supposed “donor species”. We also show that “exchanged” molecules share only 75% sequence homology, a degree of divergence incompatible with established mechanisms of recombination and genomic features of *Adineta*. These findings are parsimoniously explained by cross-contamination of tubes with animals or DNA from different species. Given the proportion of tubes contaminated in this way, we show by calculation that evidence for "intraspecific horizontal exchange" in the same dataset is explained by contamination with conspecific DNA. On the clear evidence of these analyses, the 2016 study provides no reliable support for the hypothesis of horizontal genetic transfer between or within these bdelloid species.

## Introduction

The maintenance of sexual reproduction is a fundamental evolutionary problem (Maynard Smith 1978). Despite many costs (Lehtonen et al. 2012), sex is nearly universal among eukaryotes (Speijer et al. 2015), whereas obligately asexual lineages are rare and typically short-lived (Bell 1982; Burt 2000). Various genetic and ecological hypotheses have been proposed to explain this paradox (Kondrashov 1993; Hartfield & Keightley 2012) but definitive tests are lacking, in part because of difficulties identifying appropriate study systems (Lehtonen et al. 2012; Meirmans et al. 2012).

One approach is to investigate groups that seem to have evolved without sex over extended timescales (Judson & Normark 1996; Normark et al. 2003). Whatever mechanism typically suppresses asexuality ought to be absent or unusually mitigated in these exceptional groups, whose genetics and ecology may thus illuminate the rules that maintain sex elsewhere. However, the remarkable claim of longstanding asexuality must first be established rigorously. Only a handful of 'ancient asexual' candidates have been identified, and some are subject to ongoing doubt (Lunt 2008; Schurko et al. 2009), especially when extant populations continue to invest in males (Palmer & Norton 1991; Smith et al. 2006; Schwander et al. 2013).

Rotifers of the Class Bdelloidea have attracted particular attention in discussions of asexuality. These microscopic filter-feeding invertebrates live in nearly every freshwater habitat worldwide, however tiny or ephemeral, thanks to their tolerance for extreme conditions, including complete desiccation (Ricci 1987). The main indication of longstanding asexuality is the absence of evidence for males in nearly 500 described species (Segers 2007) despite centuries of observation (Hudson & Gosse 1886; Mark Welch et al. 2009; Birky 2010). Cytological evidence is also suggestive: *Philodina roseola* has 13 chromosomes, including three without apparent morphological homologs, and oogenesis does not seem to involve reduction or synapsis (Hsu 1956; Mark Welch & Meselson 1998).

Several unusual features of bdelloid genomes have been considered as possible consequences of longstanding asexuality. Bdelloids have far fewer vertically transmitted retrotransposons than other animals (Arkhipova & Meselson 2000; Flot et al. 2013), as predicted by selfish DNA theory when sex is absent (Hickey 1982). Deeply divergent gene copies in certain bdelloid species were initially interpreted as former alleles on previously paired chromosomes that had been evolving independently for millions of years without recombination (Birky 1996; Mark Welch & Meselson 2000; Pouchkina-Stantcheva et al. 2007). However, later work clarified that bdelloids are ancestrally tetraploid, with most genes forming two divergent pairs of nearly identical sequences (Hur et al. 2009; Mark Welch et al. 2009). By convention, a typical gene has a closely related 'homolog', and a pair of distantly related 'ohnologs' (Flot et al. 2013). Each homologous pair diverges deeply from its ohnologs, and the two lineages seem too distant for genetic exchange (e.g. homology is 86% for ohnologs of the heat-shock gene *hsp82* in *P. roseola)*. Within pairs, homologous copies are similar or identical, which implies some ongoing homogenising mechanism whose nature is as yet unclear. It has been speculated that homologs serve as reciprocal templates for the repair of DNA doublestrand breaks (DSB) induced by desiccation, and that break-induced replication, recombination and gene conversion might mediate concerted evolution between them (Gladyshev & Meselson 2008; Mark Welch et al. 2008; Flot et al. 2013; Hespeels et al. 2014). These structural features and hypothetical mechanisms are central to current models of bdelloid genome evolution.

Bdelloid genomes have a very high proportion of genes of non-metazoan origin, which indicates exceptional rates of horizontal gene transfer (HGT) from foreign sources (Gladyshev et al. 2008). Initial claims of this kind in tardigrades (Boothby et al. 2015) have been rejected as artefacts of contamination after more careful analysis (Delmont & Eren 2016; Koutsovoulos et al. 2016; Richards & Monier 2016). In bdelloids, however, extensive inter-kingdom HGT is consistently supported by independent lines of evidence in multiple species, including datasets and analyses that exclude contamination as an explanation (Gladyshev et al. 2008; Boschetti et al. 2012; Flot et al. 2013; Crisp et al. 2015; Eyres et al. 2015). Most foreign genes appear bacterial in origin, but fungal, plant and archaeal genes also are represented (Boschetti et al. 2012). Animal genes have probably been acquired horizontally too, but these are not readily distinguished against a metazoan background.

How and why bdelloids have accumulated so many foreign genes is unclear. One hypothesis is that environmental DNA may be incorporated during recovery from desiccation, if cell membranes are compromised and genomes require extensive repair (Gladyshev et al. 2008; Hespeels et al. 2014). However, at least one desiccation-tolerant tardigrade shows no such accumulation (Hashimoto et al. 2016), whereas some desiccation-intolerant bdelloids continue to acquire foreign genes (Eyres et al. 2015). Although the rate of stable incorporation is high relative to other animals, it is estimated to be low in absolute terms: on the order of ten events per million years (Eyres et al. 2015).

When first describing non-homologous HGT into bdelloids from other Kingdoms, Gladyshev et al. (2008) speculated that "there may also be homologous replacement by DNA segments released from related individuals", in which case "bdelloid rotifers may experience genetic exchange resembling that in sexual populations". This idea was extended by Flot et al. (2013), who speculated that HGT “from rotifer to rotifer” may have “replaced” functions otherwise performed by sex. Here, we discuss a study appearing in *Current Biology*, which claims to support this view by supplying evidence of extensive transfer of genetic material between individual bdelloid rotifers.

Debortoli et al. (2016) collected lichen and soil from five trees in a small area and looked for bdelloid rotifers of the cosmopolitan genus *Adineta* (Hudson & Gosse 1886). Animals were moved to 576 tubes, from which DNA was extracted. PCR was used to amplify a 0.6kb region of mitochondrial cytochrome oxidase I (mtCO1), a common molecular barcode (Hebert et al. 2003). From these sequences, the authors delineated six molecular taxa (Pons et al. 2006), which they call "*Adineta vaga* Species A-F" (we follow this terminology for convenience). A subset of 82 samples representing a range of mtCO1 haplotypes from these “cryptic species” was subjected to whole-genome amplification. Four nuclear marker loci were then further amplified by PCR and sequenced.

Most samples yielded sequences characteristic of a single species at all five loci. However, for six samples (7.3%), sequences at different loci matched two or even three different species. The authors interpret this incongruence as "strong evidence" of "multiple cases" of "interspecific horizontal genetic transfers" from "donor species" to "recipient individuals." In other samples from the same dataset, the authors report “allele sharing” within “haplotype trios” of conspecific individuals, which they interpret as evidence of “intraspecific horizontal exchange”. They conclude that they have discovered "an unexpected (and possibly unique)…ameiotic strategy of genetic exchange and recombination among asexual, morphologically female organisms", which they "propose here to call ‘sapphomixis’ (from the name of the Greek lesbian poetess Sappho and mixis 'mingling')."

The dataset features some surprising patterns that seem to require clarification. In every case of reported incongruence, the inferred donor species was recovered from the same maple or plane tree as the putative recipient at the time of sampling (Debortoli et al. 2016; Table S3). This is quite unexpected, since the genus *Adineta* subsumes vast cryptic diversity. *A. vaga* alone comprises at least 36 independently evolving entities (Fontaneto et al. 2011; Robeson et al. 2011). Rotifers of this genus have high dispersal potential (Fontaneto et al. 2008; Wilson & Sherman 2013) and the lifespan of a patch of lichen is short in evolutionary terms. If genetic exchange occurs so promiscuously among such diverse and mobile animals, it is remarkable that every donor species happened to be sampled in the same small area as the recipients at the same time. Even more striking, every case of incongruence involved a sequence that was also found natively in one of the other 81 rotifers sampled, enabling the authors to construct a perfectly self-contained circular representation of the “interspecific transfers” (their Figure 4).

We hypothesised that evidence interpreted as horizontal genetic exchanges might instead result from accidental cross-contamination of rotifers or rotifer DNA between tubes during preparation of samples. This would explain the hermetically self-referencing pattern of incongruent sequences in samples showing “interspecific DNA transfers”. Specifically, more than one animal (or loose DNA from additional animals) may occasionally have been added to a single tube during the attempted isolation of 576 individuals. It is technically challenging to isolate *Adineta* individually to Eppendorf tubes, as we have found from personal experience (Supplemental Material). The animals are small even when fully extended; when disturbed, they contract rapidly into tiny, motionless, transparent spheroids that stick tenaciously to plasticware, and have about the same refractive index. If the isolation protocol used by Debortoli et al. (2016) resulted in cross-contamination of a subset of samples presented as individuals, it would give the misleading appearance of inter-and intraspecific genetic exchanges. Here, we test this alternative hypothesis using two independent and complementary sources of evidence: original Sanger sequencing chromatograms provided by the authors, and alignments of genetic and genomic data from public repositories.

## Results and Discussion

### Experimental determination of the effects of cross-contamination on Sanger chromatograms

If samples showing "interspecific horizontal genetic transfers" contained DNA from two or more animals of different species, additional haplotypes ought to be evident in chromatograms produced by Sanger sequencing of PCR amplicons. In particular, prior sequencing of thousands of rotifers indicates that each individual has only one mtCO1 haplotype (e.g. Fontaneto et al. 2009), therefore to find double peaks in mtCO1 chromatograms would provide evidence for contamination.

We conducted an experiment to determine the pattern of mtCO1 chromatogram peaks when two animals are present in one tube. We chose two bdelloid clones from our cultures: '*A. vaga* (AD008)', which supplied the reference genome for *A. vaga* (Flot et al. 2013), and '*Adineta* sp. (AD006)'. We prepared replicate Eppendorf tubes, either with a single individual, or deliberately contaminated with two individuals, one from each species (Table S1). We extracted DNA and amplified the mtCO1 marker using the methods and primers described by Debortoli et al. (2016). Bidirectional chromatograms were generated by direct Sanger sequencing with the PCR primers.

The phred quality scores of the chromatogram files (Ewing & Green 1998) were only slightly and not significantly lower for tubes with two animals versus one (mean Q20: 90% vs. 92%, N = 38, t = 1.13, P = 0.26). Even when two animals were present, the vast majority of base calls (97.9-99.6%) matched a single species (Table S2). The additional animal did not manifest as obvious double peaks at the expected polymorphic sites, but as small, subtle minority peaks, typically hidden in baseline noise (Kronick 1997) and sometimes missing entirely. Perhaps this is unsurprising: double peaks are seldom equal in height even when amplifying alleles from diploid heterozygotes (Kronick 1997). PCR is exponential, and if two animals contribute divergent templates, large biases in final amplicon representation may arise from small initial differences in numbers of cells or mitochondria, or the efficiency of lysis, DNA extraction, primer binding, denaturation, etc. (Mullis et al. 1994).

We developed a simple quantitative method, called ConTAMPR (Contingency Table Analysis of Minority Peak Ranks), to test whether the identity of the additional animal in deliberately contaminated samples could be recovered from the pattern of minority peaks (Supplemental Material). Briefly, chromatograms were aligned bidirectionally with the known sequences of both *Adineta* clones. At each site where the two reference sequences differed, we manually ranked the heights of the fluorescence trace lines corresponding to the three minority nucleotide bases, and recorded the rank for the base fitting the known contaminant sequence. Using contingency table analysis, we tested whether the distribution of peak height ranks differed from the expectation if chromatogram noise were random. For additional rigor, we measured the distribution of minority peaks fitted to a control species (A. *ricciae*, Segers & Shiel 2005) that was not present in any sample, but whose sequence identity to both AD006 and AD008 at mtCO1 was equal (87.5%).

In uncontaminated single-animal samples with *A. sp*. (AD006), rank distributions of minority peaks matching *A. vaga* (AD008) and *A. ricciae* did not differ significantly from the null expectation or from each other (Figure 1A; χ^2^ = 3.32, d.f. = 4, P = 0.51). In deliberately contaminated samples, however, minority peaks fitted the known contaminant species significantly better than the null expectation (Figure 1B; χ^2^=174.54, d.f. =2, P < 2.2 x 10^−16^) or the control species (χ^2^=22.54, d.f. =2, P = 1.27 x 10^−5^). ConTAMPR correctly detected that tubes contained two animals, identified the second sequence and differentiated it from other candidates. Multiple biological and technical replicates yielded similar results (Figure S1). This method works because PCR amplicons were directly sequenced, leaving visual evidence of minority sequences. An approach based on cloning might miss minority sequences entirely (Mullis et al. 1994). The method is therefore suitable to test the directly sequenced amplicons of Debortoli et al. (2016) for contamination by multiple animals.

**Figure 1.**
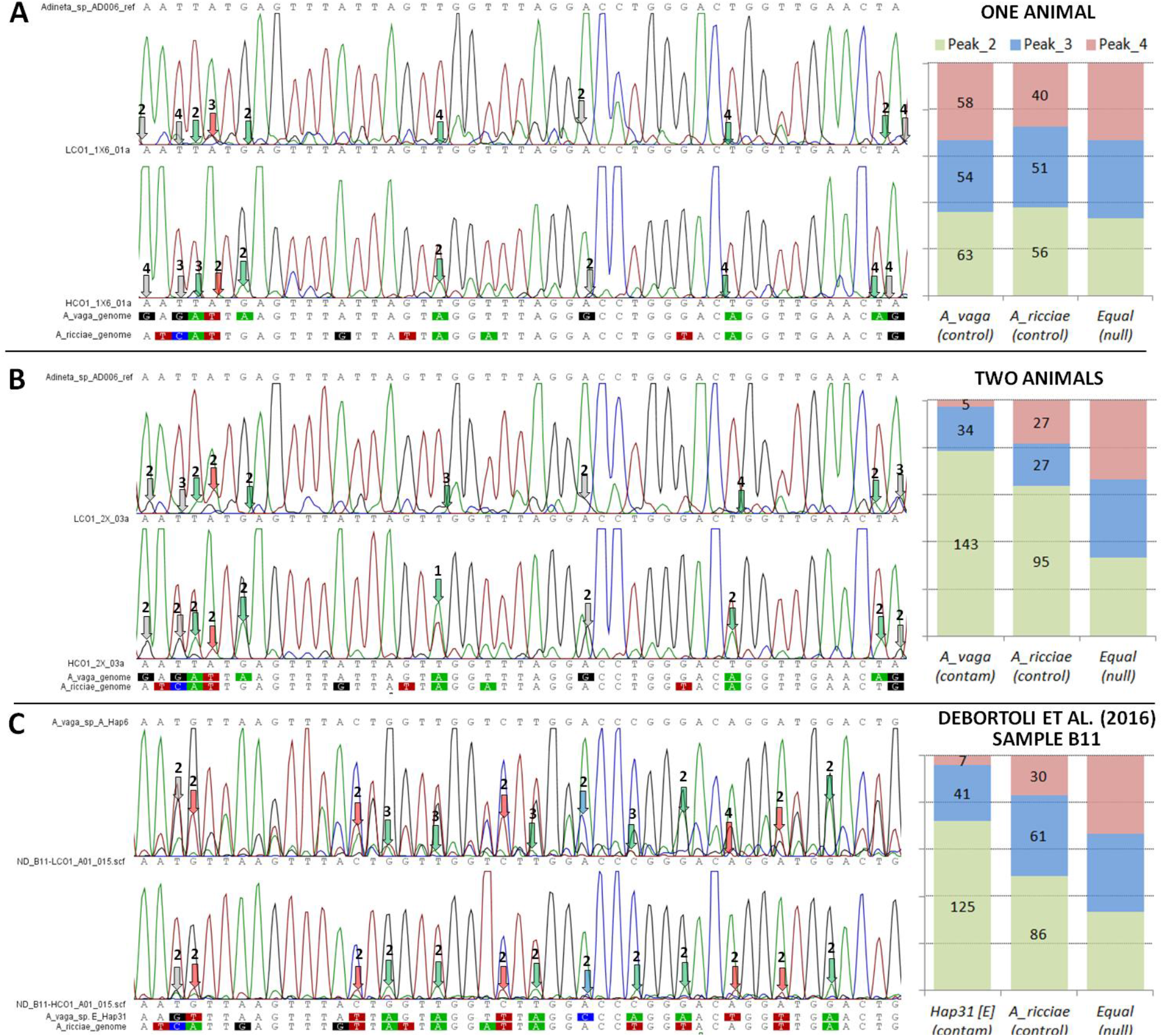
Contingency table analysis of minority peak height ranks (ConTAMPR), illustrated for mtCO1 chromatograms. **A.** For a sample with a single animal, the height ranks of minority peaks matching other rotifer sequences did not differ significantly from an equal distribution (χ^2^ = 3.32, d.f. = 4, P = 0.51). **B.** When two animals were deliberately added to a sample tube, the minority peaks were a significantly better fit to the known sequence of the second animal *(Adineta vaga)* than either the null expectation (χ^2^=88.15, d.f. =2, P <2.2 x 10^−16^), or a control rotifer sequence (A. *ricciae;* χ^2^=22.54, d.f. =2, P = 1.27 x 10^−5^). **C.** For Sample B11 of Debortoli et al. (2016), minority peaks were a significantly better fit to the predicted sequence of a suspected second animal (A. *vaga* Species E Hap31) than either the null expectation (χ^2^=127.95, d.f. =2, P < 2.2 x 10^−16^), or *A. ricciae* (χ^2^=25.39, d.f. =, P = 3.07 x 10^−6^). Minority peaks matching the focal query sequence are pointed out for a short illustrative region; numbers indicate peak height ranks. Bar graphs show the total peak height rank distributions for each sequence across the full length of the aligned locus (~605bp), and the null expectation if the peaks represent random noise.

### Evidence of cross-contamination in chromatograms generated by Debortoli et al. (2016)

To test for the signatures described above, we requested raw chromatogram files from Debortoli and colleagues. We obtained 36 out of at least 1152 mtCO1 files, including those for the six samples where interspecific transfers were claimed. We received 483 out of at least 656 chromatograms for 28S ribosomal DNA, and 133 out of at least 158 files for the EPIC25 (exon-primed intron-crossing) nuclear marker. We are grateful to the authors for their open and collaborative sharing of data.

Sample B11 was the first alphabetically to show "interspecific recombination". It was interpreted by Debortoli et al. as a single animal belonging to *A. vaga* Species A, with one mtCO1 haplotype (Hap6 [A]) and one 28S ribosomal sequence (Hap1 [A]). Other nuclear loci had supposedly acquired DNA horizontally from Species E (EPIC25 Hap35, EPIC63 Hap16 and Nu1054 Hap22). As discussed above, all three “transferred” haplotypes also were seen 'natively' in Species E animals sampled during the study. Indeed, Table S3 of Debortoli et al. (2016) shows that they all occurred together in a single animal: Individual 81 [E]. Contamination of Sample B11 with a second animal similar to this would explain the apparently incongruent signal. If so, the mtCO1 chromatograms for Sample B11 ought to show evidence of a second sequence comparable to Hap31 [E], which is found in Individual 81.

We tested this prediction using ConTAMPR (Figure 1C). The fit of minority peaks to Hap31 [E] was significantly better than the null expectation (χ^2^=127.95, d.f. = 2, P < 2.2 x 10^−16^) or the fit to *A. ricciae* (χ^2^=25.39, d.f. = 2, P = 3.07 x 10^−6^). We attempted to match the minority peaks to multiple control sequences, including the *A. vaga* reference genome and the other species delimited by Debortoli et al. (B, C, D, F). Hap31 [E] remained the best fit by far (Figure 2, χ^2^= 39.2, d.f. = 12, P = 9.73 x 10^−5^). Like other animals, single rotifers are not known to have two substantially divergent mtCO1 haplotypes. This evidence suggests that Sample B11 contained a second animal belonging to Species E, very similar to Individual 81 [E]. A parsimonious inference is that this rotifer supplied the incongruent Species E sequences at the nuclear loci where interspecific genetic exchanges were claimed.

**Figure 2.**
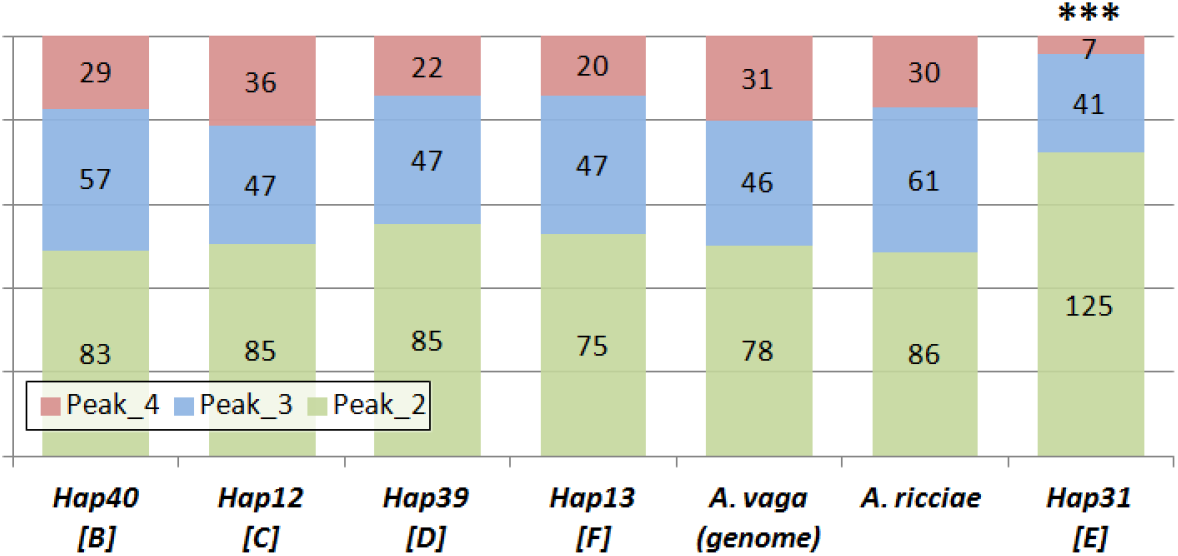
The mtCO1 chromatograms of Debortoli et al. (2016) for Sample B11 ("Individual 21" [A]) indicate a second animal belonging to Species E. The minority peaks fit Hap31[E] significantly better than six control sequences (***: χ^2^=39.2, d.f. =12, P = 9.73 x 10^−5^), which represent two reference clones and the other four species reported by Debortoli et al. (2016), and do not have significantly different distributions from each other (χ^2^=6.66, d.f. = 10, P = 0.76).

Although Debortoli et al. amplified mtCO1 directly from genomic DNA, they performed PCR for the 28S, EPIC25, EPIC63 and Nu1054 nuclear markers after whole-genome amplification (WGA) of the samples via an Illustra GenomiPhi V2 DNA Amplification Kit (GE Healthcare, Amersham Biosciences). When DNA from two animals was present, the two mtCO1 haplotypes were represented very unequally among our PCR amplicons (Figure 1B). Interposing another nonlinear amplification step introduces further opportunities for bias: some sequences might be dropped entirely, or a majority of amplicons might be generated from a minority template. This particular kit shows a bias in favor of templates with lower guanine-cytosine (GC) content (Han et al. 2012), which we suggest as an important factor in selective loss of alleles (Supplemental Material). For these reasons, it was less clear whether nuclear loci would show corresponding evidence of a second animal. However, we attempted to conduct the same analysis, first at the ribosomal 28S marker. The putative second animal was predicted to show a 28S sequence characteristic of Species E.

As predicted, the fit of minority peaks to Species E was significantly better than the null expectation (Figure 3), and also superior to Species F (χ^2^=22.45, d.f. = 2, P = 1.3 x 10^−5^), B (χ^2^=12.75, d.f. = 2, P =0. 0017), C (χ^2^=12.24, d.f. = 2, P = 0.0022) or *A. ricciae* (χ^2^ = 11.13, d.f. = 2, P = 0.0038). These results remain significant even after a conservative Bonferroni correction (α =0.0083). Species E was a better fit than Species D and the *A. vaga* reference genome too, but not significantly so, as the three species are nearly identical at the conserved 28S marker. The fit to minority peaks recapitulates a phylogenetic tree based on 28S (Figure 3): species more closely related to E fit better. The evidence at 28S therefore reinforces the mtCO1 results, indicating a second animal belonging to Species E. Because Debortoli et al. already reported Species E alleles at EPIC25, EPIC63 and Nu1054, evidence from all five sequenced loci is now brought into congruence, as predicted if Sample B11 contained an animal belonging to Species E, with a similar genotype to Individual 81.

**Figure 3.**
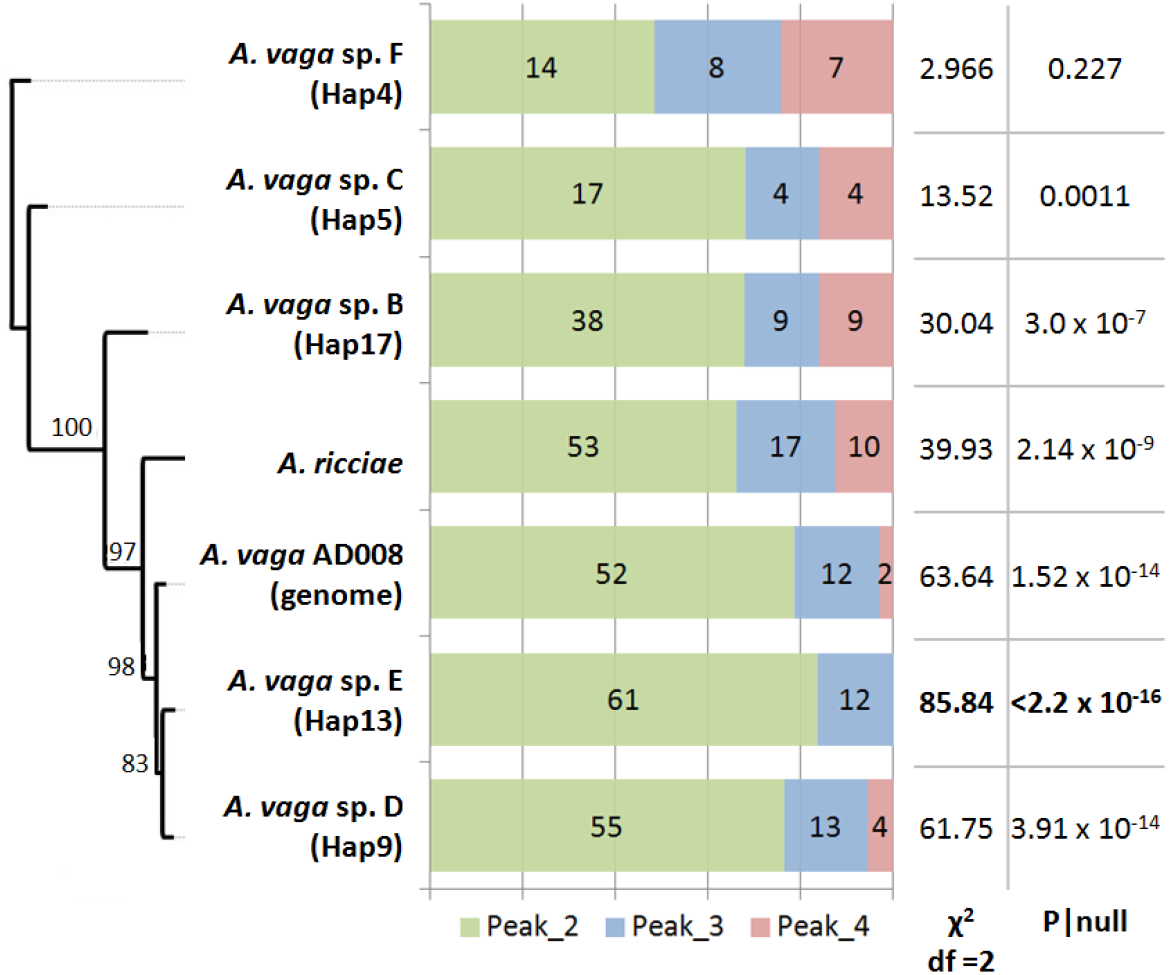
Minority peaks in 28S chromatograms for Sample B11 of Debortoli et al. (2016) indicate a second animal belonging to Species E. The probability of obtaining a peak rank distribution this extreme given random noise is lowest for Species E and increases for more distantly related control species, according to a neighbor-joining phylogeny of the 28S marker. When species distributions are compared to each other rather than to the null expectation, Species E fits significantly better than Species F (χ^2^=13.61, d.f. = 2, P = 0.0011), B (χ^2^=12.75, d.f. = 2, P = 0.0017), C (χ^2^=12.24, d.f. = 2, P = 0.0022) or *A. ricciae* (χ^2^=11.13, d.f. = 2, P = 0.0038). Species D and *A. vaga* (genome) are very closely related to E at this conserved locus, and do not have significantly different distributions.

### Evidence for "interspecific genetic exchanges" is explained by heterospecific cross-contamination

We used the same methods to investigate the other five samples where Debortoli et al. diagnosed "interspecific horizontal genetic transfers" based on incongruent species assignment among loci. Where the relevant chromatograms were provided, we aligned them bidirectionally against the sequences reported by Debortoli et al., then analysed the pattern of minority peaks with reference to the sequences of other species reported in the study (Supplemental Material).

All six samples showed significant evidence of additional sequences that were not reported by Debortoli et al. (Table 1). Three samples (B11, B22, B39) each contained two clearly identifiable mtCO1 haplotypes, which is only consistent with DNA from two animals. Even if mtDNA itself could undergo interspecific horizontal transfer (as Debortoli et al. imply for Sample B39), intergenerational bottlenecking of mitogenomes (Mishra & Chan 2014) would still result in a single mtCO1 haplotype (J.-F. Flot, pers. comm). Samples B14 and B3B1 showed evidence of DNA originating in three different rotifers (from species A, C and E). The mtCO1 chromatograms for these samples were extreme outliers in phred quality scores (Figure 4), with too much noise to narrow down just one candidate contaminant (Table 1; Figure S7).

**Figure 4.**
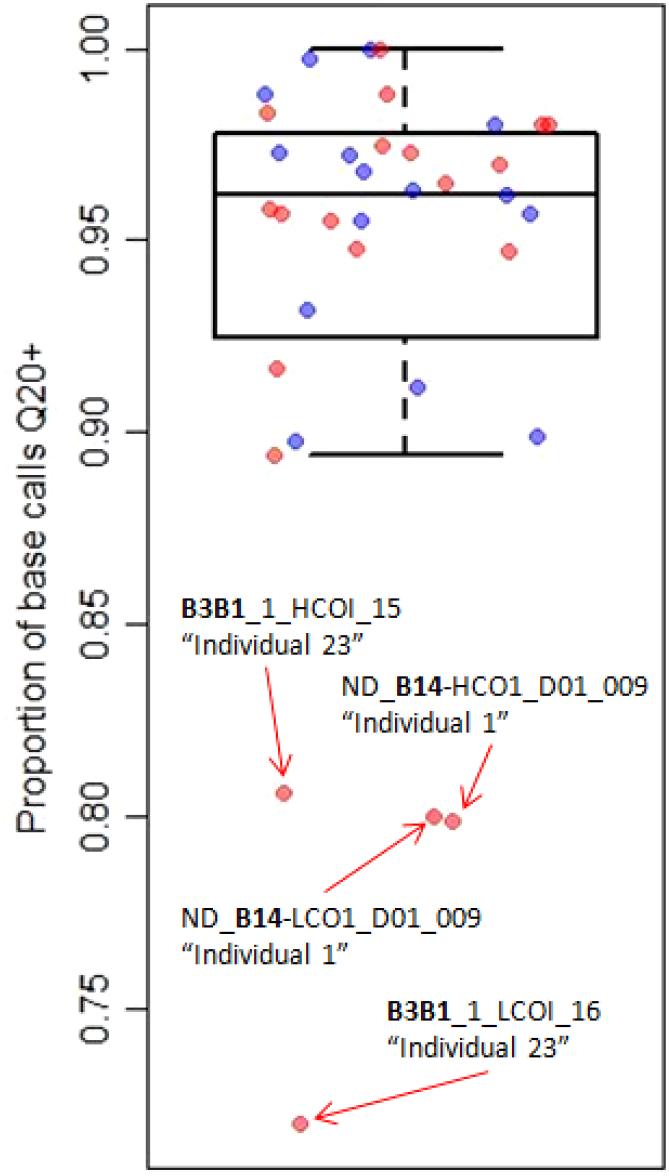
Phred quality scores for the subset of mtCO1 chromatograms provided by Debortoli et al. Red points represent samples where interspecific recombination was claimed; blue points represent samples where no such claim was made. The two groups do not have significantly different distributions (Mann-Whitney Test, N = 35, W = 127.5, P = 0.46), which matches the outcome we saw in experimental samples deliberately contaminated with two animals. However, extreme outliers in quality correspond to the only two samples with evidence of nuclear sequences from three different species (Table 1). We suggest DNA from three animals was present.

**Table 1.**
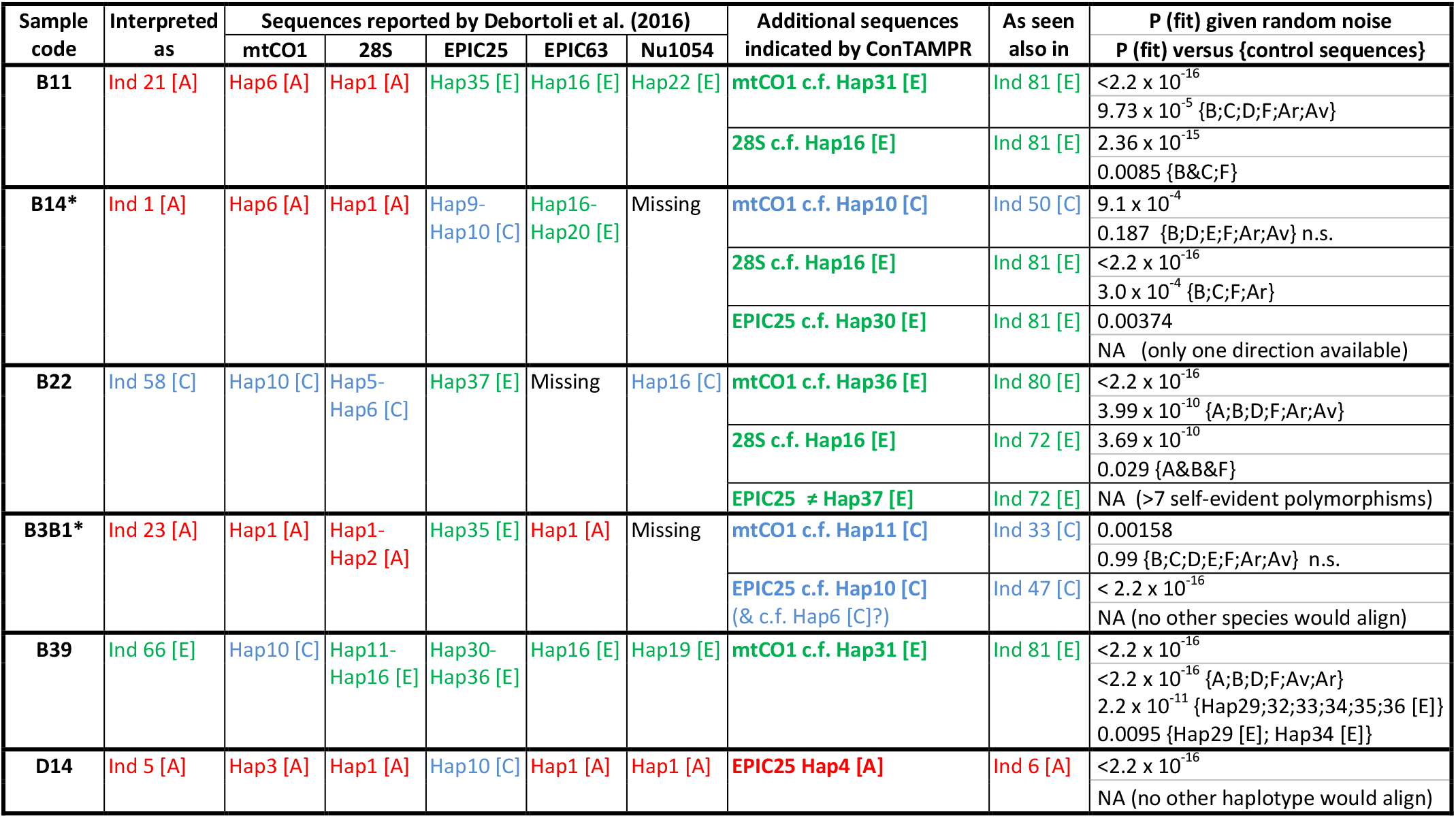
Summary of six samples with incongruent species assignments among marker loci, which Debortoli et al. (2016) interpreted as *Adineta* individuals that had acquired DNA horizontally from a “donor species”. For all six samples, minority peaks in chromatograms provided by the authors revealed additional sequences closely comparable to those seen in other animals in the dataset. These either matched the putative donor species, or the ‘native’ sequence that was supposed to have been replaced, indicating that incongruence was caused by cross-contamination. The height rank distributions of minority peaks corresponding to additional sequences were recorded, and we calculated the probability of obtaining a fit at least this good given random noise (final column, upper values). Where possible, we also recorded the equivalent fit for control species or sequences, and calculated the probability that the fit of the focal sequence shared the same distribution (final column, lower values). “Ar”: *A. ricciae;* “Av”: *A. vaga* (reference genome). Asterisks indicate samples where sequences from three different species were recovered from the same sample.

In samples B11, B14 and B22, additional 28S sequences were found. This is not an expected result of HGT, because ribosomal DNA undergoes concerted evolution (Liao 2000) that is believed to preclude the maintenance of substantially divergent 28S copies in a single rotifer, even if one arrived via HGT (J.-F. Flot, N. Debortoli & K. Van Doninck, pers. comm.). Ribosomal DNA has been amplified from hundreds of rotifers without finding copies that differ by more than 2-3 bases (e.g. Tang et al. 2012). Debortoli et al. found no more than three differences between 28S copies in any of 82 animals (their Figure 3A). In contrast, the two sequences we found in Samples B14 and B22 differ at over 40 positions in the first 700bp fragment alone. As a consequence of multiple peaks from additional sequences, quality scores for 28S chromatograms were significantly lower where transfer was claimed than where it was not (Figure 5; Mann-Whitney Test: N = 122, W = 373.5, P = 0.015).

**Figure 5.**
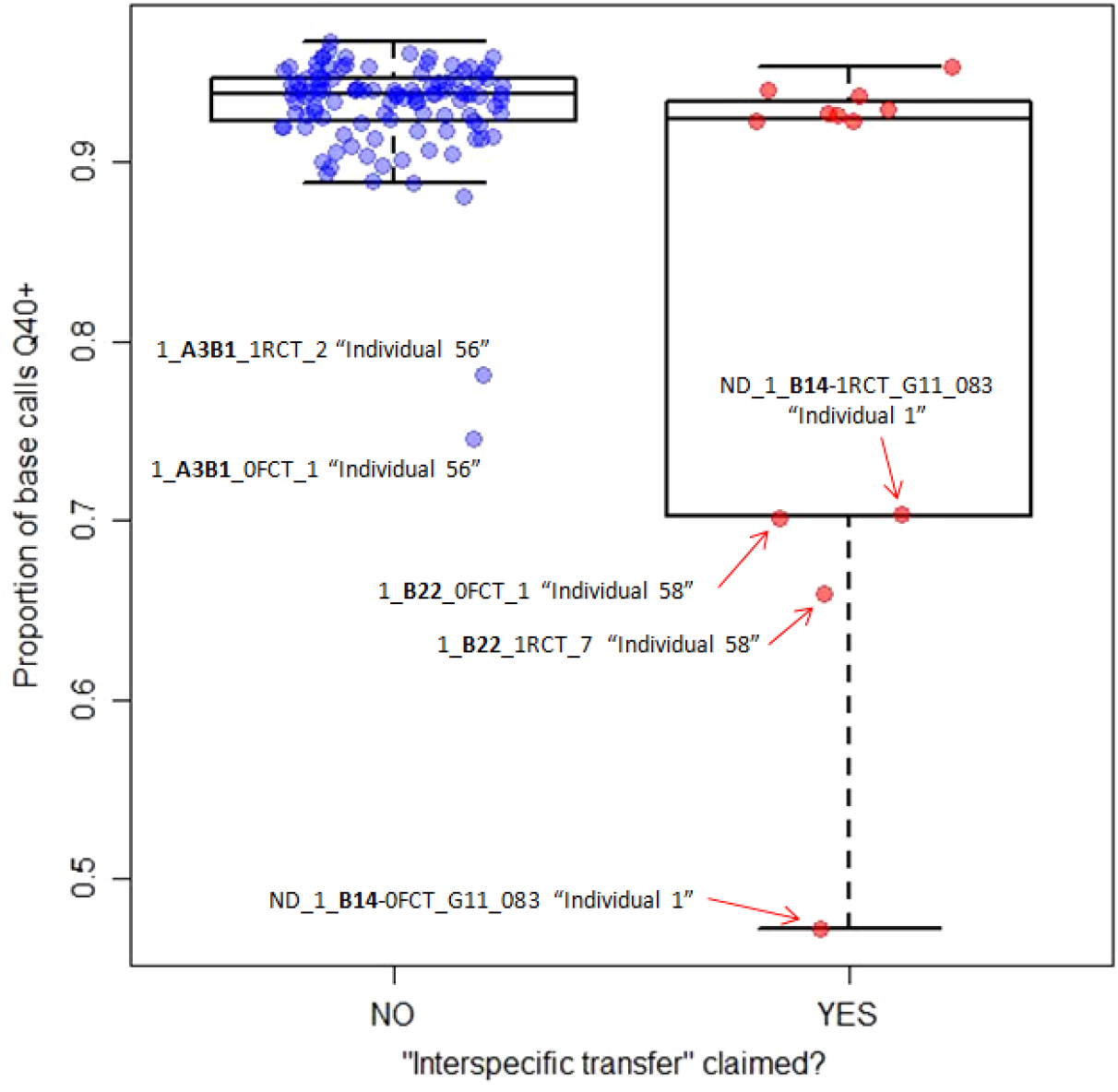
Phred quality scores for 28S chromatograms. Samples where Debortoli et al. reported “interspecific horizontal genetic transfers” have a significantly lower distribution (Mann-Whitney Test: N=122, W=373.5, P=0.015) because multiple 28S sequences are more often evident as minority peaks. This indicates that DNA from multiple animals was present in these samples, and perhaps also in some samples where “transfer” was not reported (e.g. A3B1). Sample B14 gives an extreme datapoint with a very prominent second sequence, but the two groups still differ significantly even if this is removed from the analysis (Mann-Whitney Test: N=121, W=373.5, P=0.04).

In Samples B39 and D14, we recovered the 'native' sequences that were supposed to have been replaced by horizontal acquisitions. All loci in both samples are now brought into congruence. A parsimonious inference is that they represent typical animals with haplotype associations similar to conspecifics. The incongruent sequences arise from WGA or PCR of contaminating DNA fragments, or from a second animal whose alleles were dropped during amplification at other loci. Consistent with biased or capricious amplification, the predicted 'native' sequence in Sample D14 was almost equivalent in peak strength to the heterospecific sequence after an initial PCR, but absent among amplicons from a replicate PCR using the same tube of template (Supplemental Material). If the locus had only been amplified once, there might have been no evidence of the native sequence, and the case for interspecific recombination would have been harder to reject. A lack of minority peaks after this replicate PCR confirms an important message in this dataset: even when a sample contains two different sequences, one may be dropped during any given WGA or PCR. A significant pattern of minority peaks is thus sufficient but not necessary to indicate a contaminating sequence, whereas the absence of such a pattern is necessary but not sufficient to exclude it.

### “Interspecific recombination” is not mechanistically compatible with genomic evidence

We believe mechanistic considerations independently falsify the hypothesis of interspecific recombination. According to Debortoli et al. (2016), transfers “may be mediated by DSB repair through homologous recombination (HR)". In their view, "this hypothesis is reinforced by the observation that…the transferred sequences replaced the original copies", and they "speculate that, after the integration of DNA, gene conversion promptly copied the integrated DNA on its homologous region". This last step is critical because each gene in the degenerate tetraploid genome of *A. vaga* undergoes concerted evolution with a paired homolog (Mark Welch et al. 2008; Flot et al. 2013).

In the context of HGT, Thomas and Nielsen (2005) define HR as "recombination that depends on extensive segments of high sequence similarity between two DNA molecules" (p. 714). As Debortoli et al. themselves remark, the frequency of HR "is strongly correlated with the degree of identity between the recombining DNA fragments and dramatically declines as the sequences diverge." A log-linear decline in HR with decreasing sequence identity is well established in bacteria (Watt et al. 1985; Zawadzki et al. 1995; Vulić et al. 1997; Majewski 2001; Thomas & Nielsen 2005); protists (Bell & McCulloch 2003); yeast (Datta et al. 1997); plants (Opperman et al. 2004; Li et al. 2006) and animals (LaRocque & Jasin 2010; Do & LaRocque 2015). Datta et al. (1997) found that a single mismatched base (99.7% identity) reduced HR rates fourfold in yeast, and reductions to 99% or 90% identity reduced HR rates by one and two orders of magnitude respectively. At these distances, the relationship is largely governed by active mismatch repair systems, and HR does not decline so dramatically if these are abolished. However, when sequence identity falls below 90%, rates of HR decline exponentially even in mutants devoid of mismatch repair, indicating that the machinery of HR itself fails to engage when so many bases are mispaired. At 83.5% identity, HR is effectively absent in yeast (three recombinant cells per billion: Datta et al. 1997), and rates were too low to measure even for otherwise promiscuous *Bacillus* species (Zawadzki et al. 1995).

Following this logic, Debortoli et al. remark that "closely related species should be more prone to genetic exchanges, which would explain why our study, focusing on intrageneric variation within the morphospecies *A. vaga*, detected multiple cases of genetic transfers." Indeed, an extremely high frequency of HR is required to explain their data. Even though only five loci were examined, nearly 10% of the sampled "individuals" were claimed as interspecific recombinants. To accommodate the self-contained pattern of transfers, these events must be so recent that putative donor species and recipient clones were all still living together at the time of the study, with transferred sequences remaining identical between them. Moreover, HR must have occurred at least twice in each case: once when the exogenous DNA was first integrated, and again when "gene conversion promptly copied the integrated DNA on its homologous region". Such high frequencies of homologous exchange are possible only if donor and recipient sequences have very high sequence identity.

We tested this prediction for each case of "interspecific recombination" using a simple method. We aligned each incongruent sequence from a putative donor species against the native sequence it was supposed to have replaced, and measured pairwise identities (Supplemental Material). This recreates the genetic divergences across which Debortoli et al. posit multiple, recent HR events (Table 2). The mean identity between sequences is just 74.8% (median 72.3%). This surprisingly low figure was validated for a genomic region of approximately 10kb around the 400bp EPIC25 intronic marker (Table S3, S4). Identity values are the same or lower regardless whether we consider exons, introns or intergenic regions. These numbers are not compatible with current understanding of HR in the model systems discussed above, or with current views of homologous transfer in the genome of *A. vaga*. The assembly of Flot et al. (2013) indicates that gene conversion and concerted evolution occur between sequence pairs that "are on average 96.2% identical at the nucleotide level (median = 98.6%)." Debortoli et al. (2016) claim interspecific recombination between sequences approximately an order of magnitude more divergent.

**Table 2.** Pairwise identity between DNA molecules that have undergone very recent “interspecific genetic exchange", according to Debortoli et al. (2016). This degree of sequence divergence is not compatible with current understanding of homologous recombination, or with the tetraploid structure of the *A. vaga* genome. However, it is consistent with the alternative hypothesis of cross-contamination. Divergences typically reflect substantial GC content differences among species.

| Sample | Interpreted as | Marker | Claimed replacement of DNA via HR | GC content difference (%) | Pairwise identity (%) |
| --- | --- | --- | --- | --- | --- |
| **B11** | Ind 21 [A] | Nu1054 | Hap22 [E] rpl. Hap2 [A] | -21.2 | **71.0** |
|  |  | EPIC25 | Hap35 [E] rpl. Hap1 [A] | -17.7 | **72.3** |
|  |  | EPIC63 | Hap16 [E] rpl. Hap1 [A] | -13.6 | **70.8** |
| **B14** | Ind 1 [A] | EPIC25 | Hap9&10 [C] rpl. Hap1 [A] | +4.9 | **80.2** |
|  |  | EPIC63 | Hap16&20 [E] rpl. Hap1 [A] | -13.6 | **70.4** |
| **B22** | Ind 58 [C] | EPIC25 | Hap37 [E] rpl. Hap10 [C] | -21.8 | **68.4** |
| **B39** | Ind 66 [E] | mtCO1 | Hap10 [C] rpl. Hap31[E] | -0.2 | **88.4** |
| **B3B1** | Ind 23 [A] | EPIC25 | Hap35 [E] rpl. Hap1 [A] | -17.7 | **72.3** |
| **D14** | Ind 5 [A] | EPIC25 | Hap10 [C] rpl. Hap4 [A] | +4.7 | **79.8** |
| **Mean (median):** |  |  |  | **-10.7 (-13.6)** | **74.8 (72.3)** |

The tetraploid genome of *A. vaga* supplies a second critical benchmark: each homologous pair of genes has a pair of 'ohnologous' copies whose mean identity is "73.6% (median = 75.1%)" (Flot et al. 2013). This is very close to the value we calculated for "interspecific horizontal genetic transfers", yet ohnologs have been evolving independently for millions of years within the same genomes, cells, individuals and clones, with "no conspicuous tracts of identity" between them (Hur et al. 2009). This feature of *A. vaga* militates strongly against the claim that molecules with 74.8% sequence identity could undergo frequent recombination or gene conversion within this genome, still less horizontally.

We considered the possibility that interspecific recombination might involve some alternative mechanism with less stringent identity requirements than HR. For instance, microhomology-mediated end joining involves “the use of 5-25 bp microhomologous sequences during the alignment of broken ends before joining" (McVey & Lee 2008). We measured the degree of microhomology between two "transferred" EPIC25 regions at scales from 1-40bp, but found they were no more similar than 7650 independently evolving ohnologous pairs in the same genome, in close agreement with comparisons based on global homology (Figure S9). One sequence even shared more microhomology with its own ohnolog (the EPIC63 region) than with its "transferred" partner. This excludes the hypothesis that microhomology-based mechanisms mediate interspecific HGT. If that were so, ohnologous loci could not evolve independently, as they share at least as much microhomology at every scale, and their DNA must be more abundant and accessible in any genome, cell or individual than DNA arriving across horizontal barriers.

Integration of DNA over large genetic distances must be possible at least occasionally in bdelloid rotifers, since genes with little or no homology to metazoan sequences are incorporated (Gladyshev et al. 2008). However, the mechanism in these cases would not require homologous substitution, and therefore would not prevent ohnologs evolving independently. It may involve a combination of individually unlikely events, which would explain why absolute rates appear low (Eyres et al. 2015).

### Evidence for "intraspecific horizontal exchanges" is explained by conspecific cross-contamination

Chromatogram analysis revealed that at least six samples interpreted as individuals by Debortoli et al. (2016) were contaminated with DNA from two or three animals belonging to different species. Elsewhere in the study, the authors use the same dataset to claim evidence for “intraspecific DNA exchanges”. It seems prudent to evaluate the alternative hypothesis that these cases correspond to samples that were contaminated with DNA from two animals belonging to the same species.

Debortoli et al. report three trios of “heterozygous individuals” in Species A and C that seem to share haplotypes “in a cyclic fashion…(a||b), (b| |c), and (c||a)”. The term ‘heterozygous’ is not strictly appropriate without segregation, but we follow this usage for simplicity. The authors claim that this pattern “can only be explained by recombination between individuals”, but an alternative possibility is that one or more tubes per trio were contaminated with DNA from a second conspecific animal. For example, two tubes might each contain one heterozygote: (a||b) and (b| |c), while a third contains DNA from two animals homozygous at the focal locus: (c||c) & (a||a). As the authors rightly remark, this requires no intraspecific horizontal exchange, since “the observation of three distinct haplotypes a, b, and c in heterozygous individuals with genotypes (a||b) and (b||c) may be explained by mutations and gene conversions alone”. At least six other permutations of conspecific contamination are capable of producing a haplotype trio in the same way (Table S5).

Unlike heterospecific contamination, conspecific contamination cannot be tested via chromatograms. Samples in Species A and C all share the same one or two 28S copies, and identical mtCO1 sequences are seen in animals that are homo-or heterozygous for EPIC25 haplotypes in the trios, rendering the DNA of two conspecific rotifers indistinguishable at these loci. Instead, we can apply probability theory. Evidence for inter-and intraspecific transfer relies on the same 576 tubes prepared from the same natural material. Given the relative abundances of *Adineta* species in that material, and the number of demonstrated heterospecific contamination events, we can calculate the number of tubes expected to be contaminated with DNA from animals belonging to the same species (Table S6). Even using highly conservative calculations, eight out of 82 tubes (95% CI: 4-15) are expected to be contaminated in this way. Just three such tubes would be sufficient to explain the evidence for intraspecific allele sharing; we calculate a 99.27% probability that conspecific contamination events occurred at the minimum rate required to generate three cyclic trio artefacts.

Given its high frequency, Species C is by far the most likely to show conspecific contamination. The number of tubes expected to contain DNA from two Species C rotifers is 7 (95% CI: 4-14). Debortoli et al. present genomic data for three samples interpreted as Species C individuals, selected for sequencing *post-hoc* because one PCR marker seemed to show conspecific allele-sharing. Given the calculations described above, the probability is 99.98% that one or more tubes contained DNA from two animals belonging to Species C (Supplemental Material). Contamination of this sort would also explain why the same Species C trio seemed to show “signatures of inter-individual recombination” at unlinked loci (EPIC63, *hsp82)* even though such concordance is not expected if DNA fragments at different loci undergo recombination and gene conversion frequently and independently.

The issue of transfer across non-credible genetic distances does not affect purported intraspecific recombination, since mean homology between sequences in trios is 99.1%. Instead, the opposite problem applies: when sequences differ at just 3-5 sites, “allele sharing” may need no explanation beyond “mutations and gene conversions alone", as Debortoli et al. rightly note. We tested this hypothesis for the two trios in Species A by analysing shared polymorphisms (Figure S10). In both cases, a single point mutation and a single gene conversion event are sufficient to explain the whole cycle. For these two trios, it is not even necessary to invoke the high probabilities of conspecific cross-contamination calculated above, let alone intraspecific horizontal exchanges.

## Conclusions

Debortoli et al. (2016) claimed to supply “strong evidence that inter-and intraspecific DNA exchanges occur within the bdelloid rotifer genus *Adineta*. "However, chromatograms provided by the authors reveal that samples treated as individuals were contaminated with DNA from multiple rotifer species, whose identities can be recovered statistically. The probability of these patterns emerging from random noise is vanishingly low. Accidental cross-contamination parsimoniously explains the results of Debortoli et al. without reference to extraordinary or novel genetic phenomena. We can reject the argument that some transfers are real and others artefacts, since genetic alignments show that interspecific recombination is not a biologically credible interpretation for any of the incongruent sequences reported. Pairwise identities are not compatible with homologous substitution even within the *A. vaga* genome, still less with HGT at the rate the dataset implies. Given the observed rate of heterospecific contamination, the probability exceeds 99.9% that conspecific contamination occurred sufficiently often to explain evidence for intraspecific DNA exchanges too, where any explanation is needed beyond mutations and gene conversions alone. In our view, these analyses constitute clear evidence that the findings of Debortoli et al. (2016) are unreliable. The interesting hypothesis that “bdelloid individuals of the genus *Adineta* exchange DNA within and between species” may be true or false, but this work supplies no credible evidence to address that question.

The study appears to have met with wide acceptance (Krause et al. 2016; Ram & Hadany 2016; Sharp & Otto 2016; Tilquin & Kokko 2016), perhaps because bdelloid asexuality has long been considered problematic (Maynard Smith 1986), and the work seemed to confirm the enticing idea that sex is “replaced” by HGT (Gladyshev et al. 2008; Flot et al. 2013). The high proportion of foreign genes in bdelloid genomes lends this hypothesis a superficial appeal, but current formulations are simplistic and imprecise, and a clearer theoretical framework is needed to assess its evolutionary plausibility. In particular, sex has genome-wide consequences each generation via segregation, crossing-over, independent assortment, outcrossing and sexual selection. These underpin a plethora of formal models for the maintenance of amphimixis (Kondrashov 1993). Fragmentary horizontal transfer, homologous or otherwise, is not a simple analog of meiotic sex, but has quite different population genetic consequences (Redfield 2001; Narra & Ochman 2006; Agrawal 2009; Croucher et al. 2016).

The work of Debortoli et al. (2016) was criticised by Signorovitch et al. (2016), who did not identify the issues discussed here, but were concerned that the results seemed incompatible with their own prior claim of “a striking pattern of allele sharing consistent with sexual reproduction and with meiosis of an atypical sort” in the bdelloid *Macrotrachela quadricornifera* (Signorovitch et al. 2015). As argued by Debortoli et al., “our observations do not support the hypothesis of an *Oenothera-like* meiosis” in *Adineta*, because “among the 82 *A. vaga* individuals sequenced for four nuclear markers, no trio of individuals presented congruent patterns of shared sequences” (Flot et al. 2016). This lack of evidence for congruent allele sharing cannot be attributed to cross-contamination, which would not obscure a pattern so striking and widespread as that reported by Signorovitch et al. (2015). That study itself features some arguments and patterns that seem to require empirical clarification, and we join the authors in “awaiting full genome sequencing of the allele-sharing individuals of *M. quadricornifera*” (Signorovitch et al. 2016), which will shed further light.

Schwander (2016) commented on the work of Signorovitch et al. (2015) and Debortoli et al. (2016), under the title *"The End of an Ancient Asexual Scandal”*. Schwander asserts that “these two studies show beyond doubt that genetic exchange between individuals occurs in different bdelloid species.” Schwander takes the position that “even small amounts of recombination and genetic exchange between individuals appear to be enough to provide all the benefits of sex", and therefore judges that “bdelloids should no longer be considered as asexuals.” With the advantage of hindsight, we suggest some of these rather strong assertions might now be considered premature.

In the words of Maynard Smith (1986), bdelloid rotifers “remain something of an evolutionary scandal.” Molecular inquiries have revealed some truly extraordinary features in these tiny and unassuming creatures (Arkhipova & Meselson 2000; Gladyshev et al. 2008; Mark Welch et al. 2008; Boschetti et al. 2012; Hespeels et al. 2014; Arkhipova et al. 2017), inviting speculation about links to their unusual reproductive mode. It is tempting to seek confirmation of these exciting ideas, and to expect further extraordinary discoveries. However, that approach opens the door to well-known biases, both when interpreting data and when evaluating the work of others (Nickerson 1998). Given concerns around contamination and misinterpretation, we join others in recommending that heightened scrutiny be applied to future claims of ‘non-canonical’ genetic exchange in microscopic animals (Richards & Monier 2016), and to work on the molecular genetics of bdelloid rotifers in particular. A more cautious and incremental programme will better facilitate firm progress, and help exploit the unusual leverage that bdelloids offer on fundamental evolutionary and genetic questions.

## Acknowledgements

We sincerely thank N. Debortoli, K. Van Doninck and J.-F. Flot for freely sharing chromatograms to help us test the alternative hypothesis we had proposed. This goes beyond standard practice for sharing sequencing data, which perhaps ought to be revisited. We appreciate their transparent and scholarly conduct, and their thoughtful and collegial correspondence since we first communicated our concerns in April 2016. We are grateful to M. Blaxter, G. Koutsovoulos and M. Meselson for helpful comments, and to P. MacMahon for mathematical advice. This work was funded in part by a NERC Postdoctoral Fellowship (NE/J01933X/1) to C.G.W., and NERC grant NE/M01651X/1 to T.G.B.

## Author Contributions

Conceptualization: C.G.W.; Methodology: C.G.W. and R.W.N.; Software: R.W.N.; Formal Analysis: C.G.W. and T.B.; Investigation: C.G.W.; Data Curation: C.G.W.; Writing-Original Draft: C.G.W.; Writing-Review & Editing: R.W.N., T.B. and C.G.W.; Visualization: C.G.W. and R.W.N.; Supervision: C.G.W. and T.B.; Project Administration: C.G.W. and T.B.; Funding Acquisition: C.G.W. and T.B.

